# Computational modelling of hippocampal sensitivity to expectation violation

**DOI:** 10.1101/611467

**Authors:** Darya Frank, Marcelo Montemurro, Daniela Montaldi

## Abstract

Pattern separation and completion are fundamental hippocampal computations supporting memory encoding and retrieval. However, despite extensive exploration of these processes, it remains unclear whether they are modulated by top-down processes. We used a neural network model to examine how unexpected information is represented by the hippocampus. During training the network learned a contingency between a cue and a category, which the target object belongs to. At test, we presented the network with congruous and incongruous cues, as well as perceptually similar foils. We used representational similarity analysis to examine how the top-down expectation modulation interacts with bottom-up perceptual input, in each layer. All subfields showed an interaction between the two, with DG and CA3 being more sensitive to expectation violation than CA1. A further multivariate analysis revealed that representational differences between expected and unexpected inputs were prominent for moderate to high levels of perceptual overlap in DG/CA3. This effect diminished when inputs from DG and CA3 into CA1 were lesioned. Overall, our findings suggest pattern separation in DG and CA3 underlies the effect that violation of expectation exerts on memory.

## Introduction

A fundamental goal of cognitive science is to characterise the mechanisms supporting memory formation and retrieval in the brain. One prominent model is the complementary learning systems model (CLS; McClelland, McNaughton, & O’Reilly, 1995), which posits that the hippocampus and neocortex play different roles in representing memories. According to CLS, the hippocampus supports ‘one-shot learning’, storing each experience as a unique memory representation in a process known as *pattern separation.* At retrieval, the hippocampus can reinstate the representation from a partial cue using *pattern completion.* Over time, these memories become generalised in cortex, representing commonalities of overlapping experiences. This model helps account for the apparent contradicting goals of remembering specific details, but also extracting the gist from similar events. Whilst these computations have been assessed in neuropsychological, behavioural and neuroimaging studies, a few outstanding questions remain. One of these is whether hippocampal pattern separation can be adaptively engaged to facilitate learning. Here we investigate whether top-down factors can modulate this encoding mechanism by employing a neural network model of the hippocampal subfields and a novel cognitive task.

Since Marr’s (1971) seminal work identifying the computations performed by the hippocampus, considerable evidence has indicated pattern separation and completion are vital to overcoming the catastrophic interference problem (McClelland et al., 1995; Norman & O’Reilly, 2003). To successfully discriminate between similar inputs, yet to be able to retain both of them, inputs are represented distinctly by pattern separation, but both can still be retrieved from a shared partial cue. Importantly, these processes are not mirror-images of one another, but rather are complementary; in order to reinstate (pattern-complete) the targeted memory it has to be accessible and distinct from similar experiences (pattern-separated). Therefore, the underlying biological mechanism should support their ongoing co-occurrence. Indeed, the theta cycle has been suggested to set the time-course for this synchronisation, with pattern-separation occurring at the trough and pattern completion at the peak of the cycle (Hasselmo, Bodelon, & Wyble, 2002; Ketz, Morkonda, & O’Reilly, 2013; Kunec, Hasselmo, & Kopell, 2005). The flow of information and respective firing properties of the hippocampal subfields, as recorded by electrophysiological studies, lend support to this notion. Sparse activity in DG and mossy fibre projections to CA3 are believed to underlie pattern separation, whereas the recurrent collaterals in CA3 and further projection via Schaffer collaterals to CA1 support pattern completion (Leutgeb, Leutgeb, Moser, & Moser, 2007; Yassa & Stark, 2011). The dual contribution of CA3 to separation and completion is believed to be determined by the nature of the input (Yassa & Stark, 2011). Whilst such a mechanism offers a simple yet comprehensive explanation for the bottom-up neural synchrony of pattern separation and completion, the role of potential cognitive modulators remains unclear.

One such process is top-down expectation, which has been shown to guide adaptive behaviour (Bar, 2009) and consequently modulate memory performance (Frank & Montaldi, 2019; Kafkas & Montaldi, 2018a; Long, Lee, & Kuhl, 2016). As pattern separation and completion are fundamental to memory encoding and retrieval, it is of particular interest to examine whether they drive expectation-related memory effects. Indeed it has been posited that a mismatch between predicted and received outcome prepares cellular mechanisms towards encoding novel or salient information, including a shift in the theta rhythm (Axmacher et al., 2010; Douchamps, Jeewajee, Blundell, Burgess, & Lever, 2013; Meeter, Murre, & Talamini, 2004; Mizumori, 2016). Human neuroimaging data showing hippocampal sensitivity to match/mismatches (Kumaran & Maguire, 2007; Long et al., 2016) provide further support for this notion, and together with findings from rodent studies (Vago, Bevan, & Kesner, 2007), suggest such hippocampal involvement could be mediated by dopaminergic inputs from midbrain regions (Fitch, Sahr, Eastwood, Zhou, & Yang, 2006; Kafkas & Montaldi, 2015; Lisman & Grace, 2005; Pine, Sadeh, Ben-Yakov, Dudai, & Mendelsohn, 2018; Shohamy & Wagner, 2008). Recently, we have shown top-down expectation can dynamically engage pattern separation to support memory for unexpected items, as a function of the level of similarity or overlap between inputs (Frank & Montaldi, 2019). Although the behavioural task used was heavily dependent on successful pattern separation, we did not have direct recordings or simulations of hippocampal subfield activity. Here, therefore, we use a neural network model of the hippocampus to test how memory representations in different layers of the hippocampus are modulated by a violation of a learned expectation. This approach allows us to bridge the gap between different levels of analysis, combining a cognitive task adapted from human research (Frank & Montaldi, 2019), and a neural mechanism characterised by computational and animal models (Douchamps et al., 2013; Norman & O’Reilly, 2003).

A common approach to simulate the CLS hippocampal system is a neural network model implemented in Emergent (Aisa, Mingus, & O’Reilly, 2008), which has been shown to emulate behavioural and neuropsychological findings in rodents and humans (Elfman, Aly, & Yonelinas, 2014; Norman & O’Reilly, 2003; Pilly, Howard, & Bhattacharyya, 2018). This model incorporates the neurocomputational properties of the hippocampus on a small scale (a few hundred neurons), and can be used to advance our understanding of the mechanisms supporting memory encoding and retrieval. For example, Elfman et al. (2014) used perceptually similar inputs and measured the distribution of activation in different layers in two tasks to show the network could account for both high-level perception and memory. Given this biologically-constrained architecture, we can also examine patterns of activation elicited in the network in response to different inputs to investigate how memory representations are created. To this end, a recent study tested whether the hippocampus can represent statistical regularities by using sequences of items and recording the similarity of activation patterns elicited by a higher-level structure in the data (Schapiro, Turk-Browne, Botvinick, & Norman, 2017).

Neural network models can also be used to examine how external factors modulate encoding and retrieval processes within the hippocampus. For example, Meeter et al. (2004) simulated hippocampal mode shifting in response to acetylcholine release tied to novelty detection. Another, more recent, study (Pilly et al., 2018) simulated rodent data showing top-down signals from prefrontal cortex (PFC) can reduce contextual interference in the hippocampus. These studies clearly demonstrate the important role played by computational models in explaining memory phenomena (Gluck, Meeter, & Myers, 2003) and their flexibility in simulating perceptual inputs and higher-level cognitive modulators. Whereas both the effects of perceptual similarity (Elfman et al., 2014) and contextual novelty (Meeter et al., 2004; Pilly et al., 2018) have been addressed in previous models, to the best of our knowledge, the interaction between contextual surprise and input perceptual similarity has not been studied. The model used here combines these two factors to elucidate the mechanism underlying the beneficial effect of surprise on memory.

In the current study, we employ a neural network model of the hippocampus to investigate how interactions between top-down expectation and bottom-up perceptual inputs are represented in different subfields of the hippocampus. This method allows us to directly examine whether improved memory for unexpected events is supported by pattern separation. During training, the network is presented with randomly created objects and cues (see *Stimulus presentation* and Figure 1). The cues are simulated to represent a previous item, and two objects share the same cue to reflect their semantic category (man-made or natural, see Frank & Montaldi, 2019). After the network learns these contingencies, we test the originally studied inputs, as well as parametrically-manipulated similar foils (from 85% to 33% overlap with original input), akin to perceptually similar foils from Frank, Gray, and Montaldi (2019). Additionally, each object is tested once with the original cue (expected condition) and once with the opposite category’s cue (unexpected condition). Based on our previous behavioural results, we hypothesised unexpected items would elicit more distinct (dissimilar) representations compared to expected ones, and that this pattern would be more prominent in moderate to high similarity foils. Furthermore, if pattern separation is indeed the underlying mechanism of the beneficial effect of surprise on memory, we expect to see these effects more prominently in DG and CA3. To test these hypotheses, we performed representational similarity analysis (Kriegeskorte, Mur, & Bandettini, 2008) for each object and associated foils, and compared the dissimilarity between conditions across layers.

**Figure 1.**
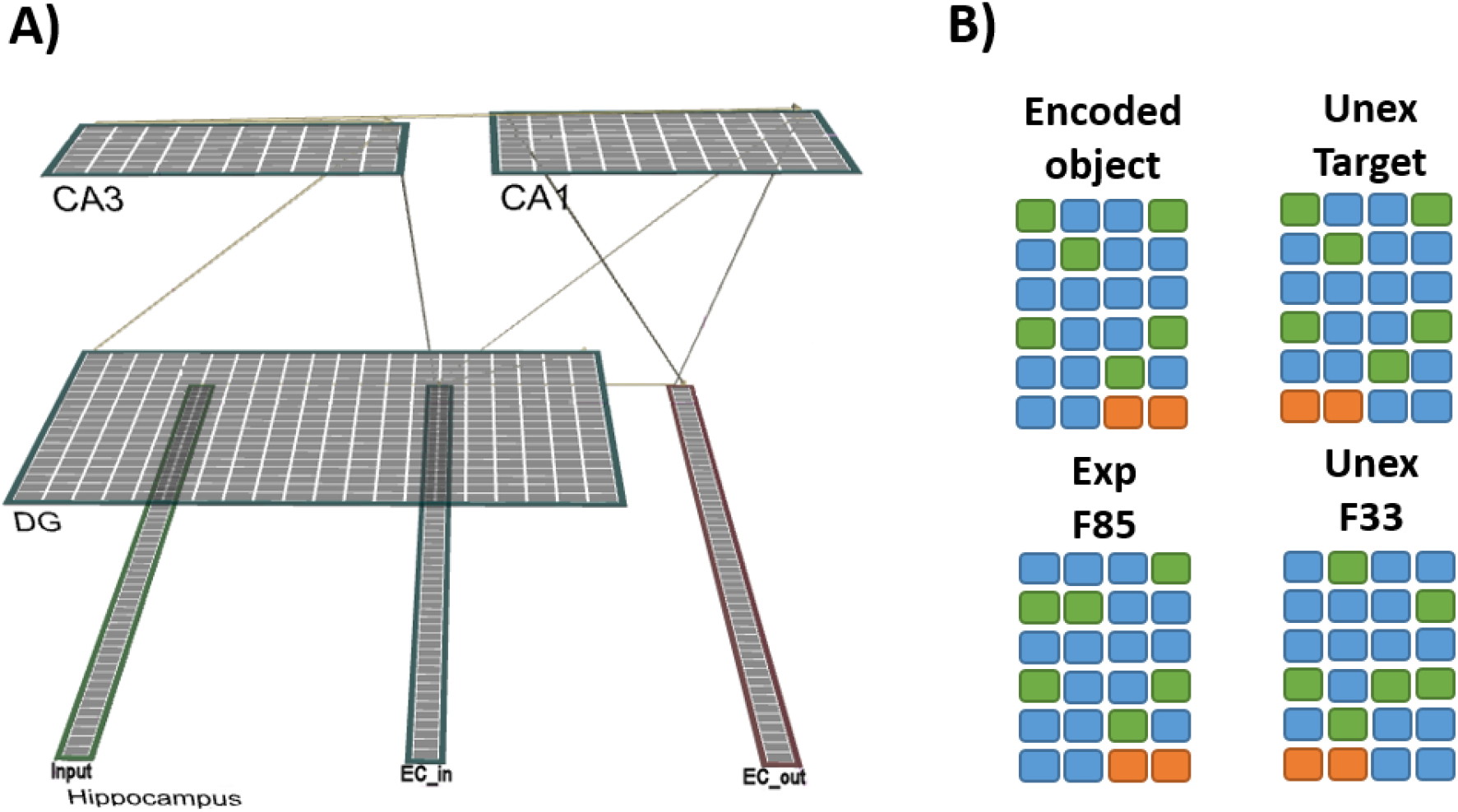
**A) Network architecture.** Illustration of the hippocampal network in Emergent. **B) Simplified representations of stimuli.** Green slots represent active stimulus units (clamped to 1) and orange slots represent the category cue (clamped at 0.9). The top left corner shows the object the network was exposed to during training. In the other corners there are examples of items presented during testing, varying in expectation condition and level of overlap.

## Results

To assert the network learned to represent the different categories used (man-made and natural), we tested whether between-category dissimilarity was greater than within-category. A 40×40 RDM was computed representing all of the trials tested, sorted by object and expectation condition. As can be seen in Figure 2, within-category dissimilarity is lower than the between-category one. To compare these differences we created four models and compared the data RDM to each of these categorical models using Kendall’s Tau A. The theoretical models represented main effects of category (man-made vs. natural), object (A vs. B vs. C vs. D), expectation (Exp vs. Unex) and a random model. We found the second-level similarity between the data RDM and the category and object RDMs was significant in all layers (all p’s < 0.01), whereas the expectation and random models could not explain the data (See Supplementary Figure 1 for second-order correlations). This pattern indicates that the overall structure of the data was most affected by the object’s own identity and the category it belonged to.

**Figure 2.**
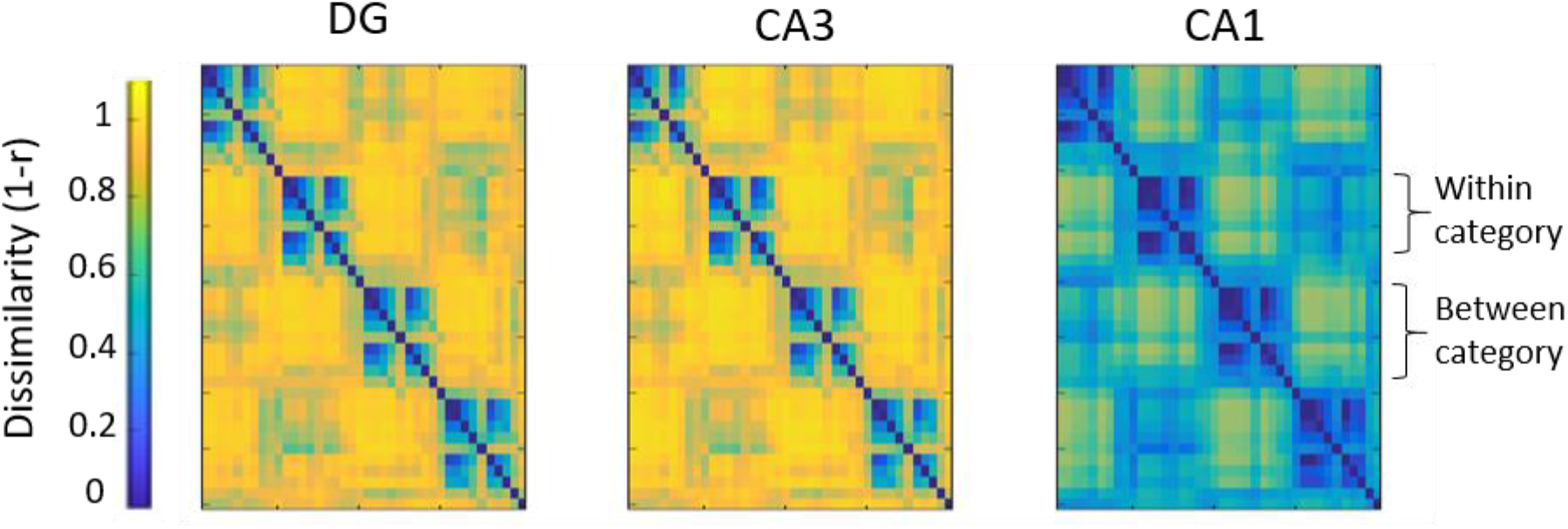
Overall 40×40 RDMs. These represent all of the test trials presented to the network and their corresponding dissimilarity (1-Pearson’s r), split by hidden layer. The four objects used are clearly represented along the diagonal, with the split in the middle of each one representing the expected and unexpected conditions tested. Objects 1 and 3 shared the same category, as did objects 2 and 4. Between-category dissimilarity is higher than within-category. Warmer colours indicate more dissimilarity.

We also tested whether the network successfully captured the amount of overlap between the trained inputs and tested foils. Sets (target and foils) from both expectation conditions were averaged across objects and compared to three theoretical models. The first model represented a scaled response, mirroring the percentage of overlap between inputs. The second model represented a thresholded response, with high overlap foils (F85 and F67) being more similar to target than F50 and F33. Finally, the third model represented a random distribution. Whilst both theoretical models outperformed the random model, using second-order Pearson’s correlations, the scaled model showed the best performance (see supplementary Figure 2). This can also be seen intuitively in the RDMs (Figure 3), with decreasing levels of overlap associated with more representational dissimilarity in all hippocampal layers. This effect was most prominent in DG and CA3 layers, consistent with their role in pattern separation.

**Figure 3.**
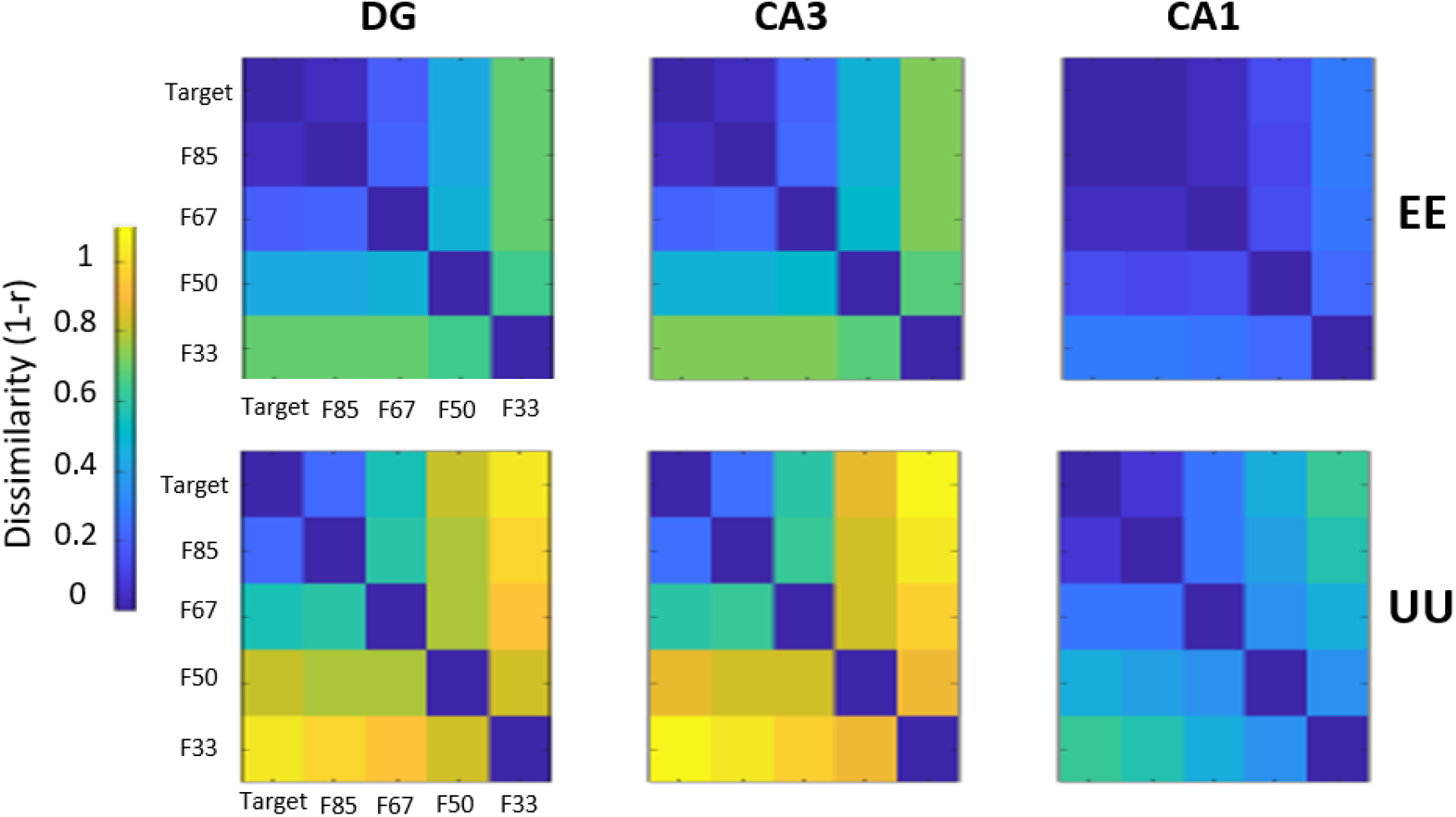
5×5 Object RDMs split by condition. The top row shows RDMs for Expected-Expected (EE) whereas the bottom row shows Unexpected-Unexpected (UU) ones. There is greater representational dissimilarity in UU compared to EE in all hidden layers.

Next, we sought to examine whether patterns of activations elicited by unexpected items differ from those elicited by expected ones. In a univariate analysis, each cell (i.e. pairwise distance) in the UU RDM was subtracted from its EE counterpart. We then subjected the data to a two-tailed t-test against a value of 0 (no difference between conditions). Across the three layers, all differences were significantly smaller than 0 (all p’s < .001) indicating larger dissimilarity in UU compared to EE RDMs. However, as can be seen in Figure 3, these differences varied in magnitude both between layers and overlap levels. In order to assess which RDM cells had the most variability between EE and UU, we turned to a multivariate approach. Singular Value Decomposition (SVD) analysis, is a dimensionality-reduction technique used to expose the substructure of a given dataset. In this analysis, a matrix or an image can be compressed whilst preserving its most informative features. Using this method we were able to capture higher-order structural differences between expected and unexpected RDMs (i.e. identifying the most informative cells in the matrix). For each layer, we used the EE-UU subtracted RDM. This RDM was then decomposed using SVD, keeping two singular values. Figure 4a depicts the reconstructed matrix, keeping 2 singular values, reflecting a compression ratio (CR) of 0.88 (warmer colours indicate no difference between conditions).

**Figure 4.**
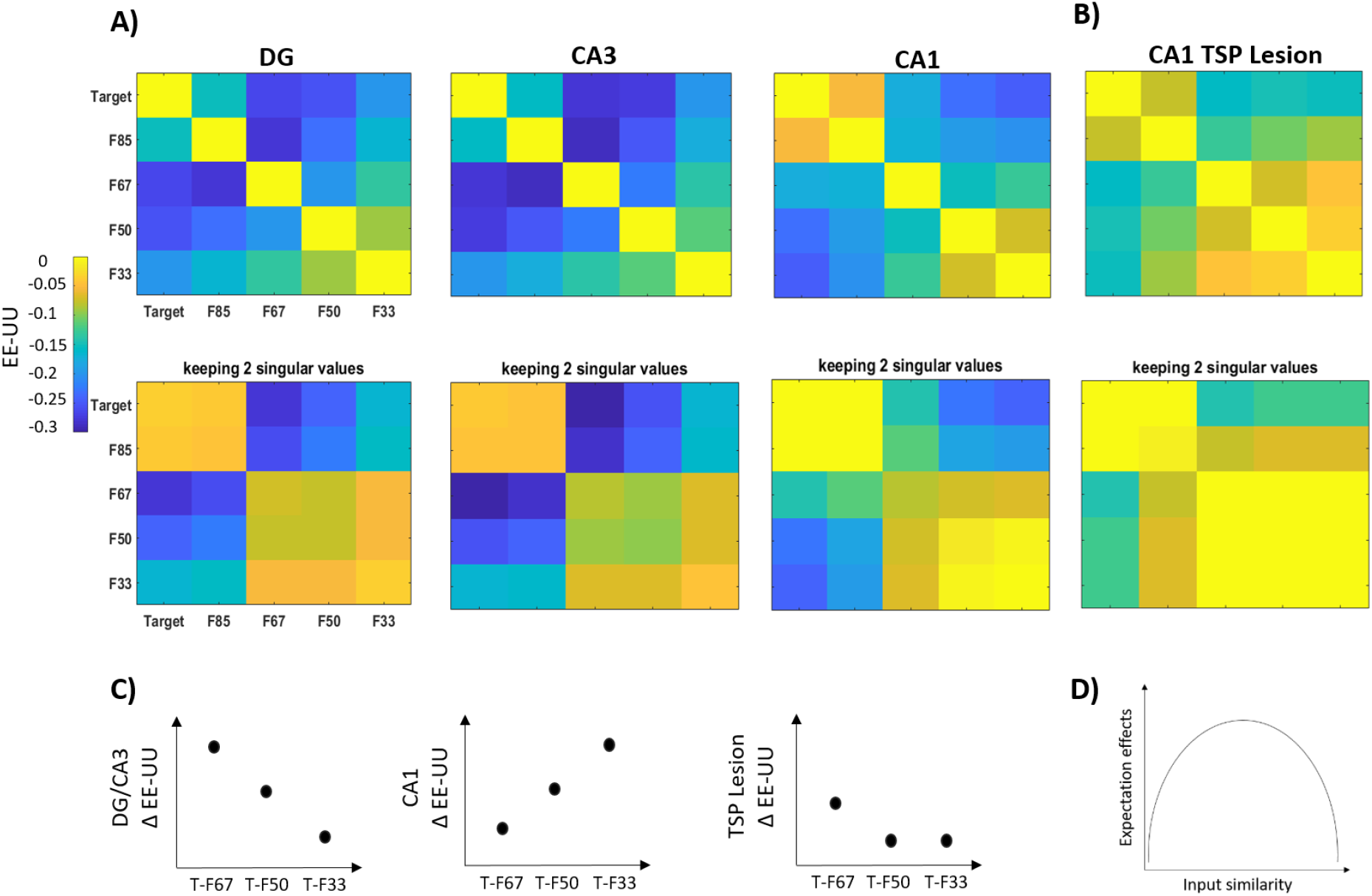
**A) SVD analysis for EE-UU RDMs.** The top row shows the raw EE-UU RDMs for each layer, the bottom row shows the SVD results, keeping the most informative two singular values. Minimal differences were observed for very high and low overlap, whereas moderate-high levels showed a graded difference between conditions. **B) SVD analysis for TSP lesion.** When inputs from DG and CA3 are muted, the observed differences between EE and UU RDMs diminish, and expand to higher levels of overlap. Warmer colours indicate less EE-UU difference. **C) Illustration of DG/CA3 and CA1 moderate effects.** Whilst DG and CA3 show a negative linear relationship between moderate to high levels of overlap and EE-UU dissimilarity, CA1 shows the opposite effect. **D) Conceptual illustration of top-down and bottom-up interaction.** Expectation effects peak at moderate to high levels of input similarity. Figure adapted from Frank and Montaldi (2019).

In DG and CA3, when target-foil similarity is very high (F85), the differences between EE and UU matrices were minimal. However, when comparing target and F67 to F33 foils, a graded difference emerged; as target-foil overlap decreases, the representational dissimilarity between expected and unexpected stimuli diminished (EE and UU are represented similarly). Finally, when mid and low similarity foils were compared to one another (as foils share little overlap amongst themselves, these represent low similarity), there were again minimal differences between the expected and unexpected conditions. This analysis suggests the representational difference between expected and unexpected RDMs in DG and CA3 was characterised by an interaction between level of overlap and expectation condition. In CA1, the SVD analysis revealed even smaller differences between EE and UU for target-F85 and between mid-low similarity foils. However, when comparing target and F67 to F33 foils, the pattern in CA1 mirrored that of DG and CA3. As overlap decreased, the difference between EE and UU increased. To ensure these higher-order structures were significantly different from arbitrary noise, we randomly shuffled each layer’s RDM 10,000 times. In each permutation we computed Spearman’s rank correlation between the shuffled and the original matrices (similar results were obtained using Pearson’s correlation). The aggregated correlations were compared to 0 (no correlation between shuffled and data matrices) using a two-sided t-test. None of these tests was significant (all p’s > 0.1), indicating the subtracted EE-UU RDMs were not significantly correlated with random noise.

Finally, we examined whether the difference between EE and UU is reduced when TSP is muted. To do so, we ran the same SVD procedure on the lesion data (only in CA1, as projections from DG and CA3 are set to 0) and compared the result to the data presented above. As can be seen in Figure 4b, whilst the overall structure in CA1 remained similar, the differences between expected and unexpected diminished (warmer colours in the lesion data). Furthermore, there are less differences observed for F85 correlations, compared to data without the lesion. To quantify the changes introduced by the TSP lesion, we compared the raw EE-UU RDM in CA1 from both datasets using a two-tailed t-test. For the T-F85 correlation difference, lesioned CA1 showed larger differences between EE and UU than the spared network t(199) = 3.251, p = .001. The T-F67 difference was significantly smaller in the lesion data t(199) = −2.02, p = .044. All other differences were in the same direction, with EE-UU differences being smaller in the lesioned TSP data (all p’s < .001).

## Discussion

Here we tested how interactions between top-down expectation and bottom-up perceptual inputs are represented in different layers of a hippocampal neural network model. We found the network learned to represent the perceptual similarity of inputs and that this representation was modulated by a violation of the learned cue-item contingency. Unexpected inputs showed greater representational dissimilarity compared to expected items. This effect was modulated by the degree of overlap between the originally encoded item and the current input, as well as the layer tested. In very high and low levels of item similarity, minimal differences were observed between expected and unexpected inputs. However, in moderate-high levels, the magnitude and direction of the effects differed between DG/CA3 and CA1. Firstly, in DG and CA3 differences between expected and unexpected inputs were greater than in CA1. Furthermore, the overall structure of this interaction differed between layers, with DG and CA3 showing a positive linear effect, whereas CA1 showed a negative one. Finally, a lesion to TSP resulted in smaller differences between EE and UU RDMs in CA1, suggesting pattern-separated inputs from DG and CA3 drive this interaction. Taken together, our results demonstrate that violation of expectation elicits an adaptive mechanism that is sensitive to the level of similarity between bottom-up inputs and existing representations.

We first asserted our network captured the bottom-up perceptual manipulation of item similarity. Indeed, the level of overlap between studied target and similar foils was reflected in the pattern of activation in each layer. In CA1, we observed higher levels of similarity, with the most similar inputs (target and F85) being undistinguishable. This is in line with CA1’s role in the integration of input commonalities (Schapiro et al., 2017). In accordance with its biological and computational pattern-separating properties (Leutgeb et al., 2007; McClelland et al., 1995) we found DG represented items most distinctly, even when the level of overlap was high. In fact, CA3 and DG exhibited almost identical patterns of activity, suggesting CA3 was more biased towards pattern separation than completion in this paradigm. It is also important to address the role of pattern separation in this context. In its most abstract sense, pattern separation consists of the automatic (blind) orthogonalisation of inputs (such that A and A’ are encoded as distinctly as A and B), but also captures the residual overlap, which ensures A and A’, which share more information than A and B, are stored ‘closer’ to each other (Hulbert & Norman, 2015). The latter computation was reflected in our data, as well as previous accounts (Schapiro et al., 2017; Yassa & Stark, 2011), showing a scaled response, with increasing representational dissimilarity as the amount of overlap decreased.

Next, we examined whether these patterns differed between expectation conditions. During training the network was taught a contingency between a cue and an object’s category. At test, expected items had the same cue they were studied with, whereas unexpected items were presented with the opposite cue. Critically, as the network was exposed to both cues during training, it is the contingency between the cue and the object that is critical for our manipulation, and not the novelty of the cue per se. Therefore, whereas foils represent a perceptual mismatch (stimulus novelty), our top-down cue manipulation is dependent on a previously learned rule. Indeed, pairwise comparisons showed unexpected items exhibited higher dissimilarity across all inputs and layers. This finding corroborates our previous behavioural results (Frank & Montaldi, submitted) showing unexpected foils were more likely to be correctly rejected than expected ones. The increased dissimilarity reported above points to the formation of a more distinct representation. This is complementary to subsequent memory effects, where targets that are successfully remembered show greater encoding-retrieval pattern similarity (Xue, 2018). In terms of the subsequent mnemonic decision, this would mean discrimination between the originally encoded targets and tested foils is easier (i.e. as the ‘distance’ between a target and a foil increases, the chance of correctly rejecting the foil increases).

We previously postulated that the advantageous effect of surprising information on memory performance is related to pattern separation (Frank & Montaldi, 2019). Specifically, we argued that the unexpected information engages pattern separation, which facilitates improved memory. To test this hypothesis, we compared the expectation conditions across the levels of overlap and layers. Although for very high (target-F85) or low overlap (foil-foil) EE and UU differences were minimal in all layers, the SVD analysis revealed that our expectation manipulation elicited distinct patterns across layers in moderate-high levels of overlap, with unexpected items showing increased dissimilarity compared to expected ones. In CA1, as target-foil overlap decreased, the representational dissimilarity between expected and unexpected items increased. This suggests the expectation violation had a more prominent effect in the lower levels of the item-similarity scale. CA1 receives pattern-separated inputs from CA3 and projections from EC, reflecting retrieval of existing representations (Norman & O’Reilly, 2003). Given these connections, CA1 has been postulated to act as a match/mismatch detector (Chen, Olsen, Preston, Glover, & Wagner, 2011; Elfman et al., 2014; Valenti, Mikus, & Klausberger, 2018). Our results suggest CA1 representations are indeed sensitive to mismatches, both perceptual and memory-based (Elfman et al., 2014), however, to different extents. The interaction between these mismatches is most pronounced at lower levels of perceptual interference, perhaps reflecting integration of items that were previously separated (T-F67) and a mismatch response for less similar ones (T-F33).

In DG and CA3, on the other hand, a mirrored effect was observed. As target-foil overlap decreased, the representational dissimilarity between expected and unexpected stimuli diminished. Whilst very high or low levels of overlap offer too much or too little perceptual interference, respectively, moderate-high overlap between encoded and current inputs require engagement of pattern separation (Norman & O’Reilly, 2003; Yassa & Stark, 2011). Within these levels of overlap, more distinct representations were observed for unexpected inputs on the higher levels of the scale. This suggests that at peak perceptual interference levels, unexpected items engage pattern separation more than expected ones. This finding dovetails with the suggestion that unexpected information can bias hippocampal computations towards an encoding state (Axmacher et al., 2010; Gruber et al., 2018; Kafkas & Montaldi, 2018b; Lisman & Grace, 2005; Shohamy & Wagner, 2008). Our findings are the first to show the consequences of such a shift, creating more distinct memory representations for unexpected items. Taken together, the contrasting results from DG/CA3 and CA1 offer an interesting view on the division of labour between these subfields when it comes to mismatches originating both from bottom-up and top-down sources. Our findings suggest top-down unexpected information is represented more distinctly in DG/CA3 when bottom-up interference is also high, whereas CA1 is more responsive to top-down manipulations when interference from bottom-up inputs is lower.

Finally, we examined how CA1 representations change when a TSP lesion is introduced, muting inputs from DG and CA3. Sparing the MSP means the resulting CA1 activation is driven by inputs from EC. Despite this lesion, CA1 representations still managed to capture the higher-order structure of the data, in accordance with previous models (Schapiro et al., 2017). Nevertheless, without the sparse inputs from TSP, differences between expected and unexpected items diminished considerably in CA1. This suggests that improved memory performance for unexpected items is driven by pattern separation in DG/CA3. It is also important to note how these findings fit with the dynamic nature of hippocampal processing. Previous research suggests pattern separation and completion occur at different stages of the theta cycle (Hasselmo et al., 2002; Kunec et al., 2005). Whilst the learning algorithm of the hippocampal model used here reflects these oscillatory properties (Ketz et al., 2013; Schapiro et al., 2017), due to the nature of the task simulated, our manipulation and tests were conducted at retrieval, where weights are not adjusted. Therefore, although our layer-by-layer analysis indicates pattern separation underlies the effects reported here, future electrophysiological research could examine the online dynamics of these effects (Axmacher, Mormann, Fernández, Elger, & Fell, 2006; Hanslmayr, Staresina, & Bowman, 2016) and how they relate to pattern completion (e.g. failure to pattern complete to target). Based on the differential pattern of responses across layers and the lesion data (most prominent effects in DG/CA3), as well as previous findings (Axmacher et al., 2010; Douchamps et al., 2013; Gruber et al., 2018; Meeter et al., 2004), our model predicts that violation of expectation would modulate hippocampal theta cycle towards encoding.

Taken together, our results offer a mechanism for the interaction between top-down expectation and bottom-up perceptual inputs and its effect on memory representations. When the level of overlap between existing and current input is moderate to high, violation of expectation helps disambiguate these representations by engaging pattern separation. However, at the two extremes, very high and low levels of overlap, this mechanism is not engaged, for different reasons. When overlap is very high, this mechanism could be turned on, and, alas, fail to exert an effect (failure of pattern separation). Conversely, when the level of overlap is low (foil-foil correlations in the current study), disambiguation is redundant, and therefore further engagement of pattern separation is unnecessary. The findings reported here have important implications for our understanding of top-down and bottom-up interactions in memory. These results reflect the dynamic nature in which memories are created and retrieved, and the sensitivity of these processes to discrepancies between current inputs and existing representations.

## Methods

### Model architecture

We used a neural network model of the hippocampus implemented in the Emergent simulation software (v.7.0.1) (Aisa et al., 2008; See Figure 1a for illustration). The model’s architecture and projections are based on the hippocampal component of the CLS framework (McClelland et al., 1995; Norman & O’Reilly, 2003)

The model includes entorhinal cortex input (EC_in_) and output (EC_out_), dentate gyrus (DG), CA3, and CA1 layers. Inputs are presented to the model via an input layer with one-to-one connections to ECin, which then projects to DG, CA3, and CA1 via the trisynaptic pathway (TSP). TSP is believed to support encoding of new memories and conjunctions, with pattern-separated representations generated in DG, and transferred to CA3 via the sparse mossy fibres (each CA3 unit receives input from 5% of DG). Recurrent collaterals in CA3 (modelled as a fully-connected projection) then allow pattern completion from partial cues to occur. The pattern-completed representation is then projected onto CA1 via the fully-connected Schaffer collateral pathway. EC is also connected directly to CA1 via the fully-connected monosynaptic pathway (MSP), which supports memory retrieval by associating direct input with diffuse inputs from the Schaffer collaterals. The network parameters of these projections are outlined in Supplementary Table 1.

Activity levels of units in the network ranged between 0 and 1. Each unit’s activity level was modulated by local inhibition from other units within the same layer, modelling inhibitory interneurons. Following previous work (O’Reilly & Munakata, 2000; Schapiro et al., 2017) this inhibition was implemented using a k-winner-takes-all equation (see Supplementary Table 2 for specific kWTA values used in each layer). In EC_in_ and EC_out_ k = 12, which is the number of active units in each stimulus presented to the network (excluding the previous cue, see *Stimulus presentation section*). Similar results were obtained using a lower inhibition at k = 15 (total number of stimulus + cue units).

The network utilises a combination of Hebbian and error-driven learning (Ketz et al., 2013; Schapiro et al., 2017) to adjust its connection weights such that an input presented to EC_in_ can be reproduced in EC_out_. To achieve this, the network goes through minus and plus phases during the learning period, akin to theta oscillations (Ketz et al., 2013). In the minus phase, EC_in_ projects to CA1 whilst CA3 input is inhibited, followed by a reversed effect whereby CA3 input to CA1 resumes while EC_in_ inputs are weakened. Theta troughs and peaks have been suggested to reflect encoding (driven by external inputs) and retrieval states (driven by internal inputs), respectively (Hasselmo et al., 2002; Kunec et al., 2005). In the plus phase, the network is exposed to the ‘ground truth’ via a loop between EC_in_, EC_out_ and CA1, during which CA1 is forced to represent the correct output (given the symmetry between EC_in_ and EC_out_). The goal of the network is to adjust its weights such that activity in the minus phase resembles activity during the plus phase, thus constantly reducing the error between minus-plus phases.

### Stimulus presentation

The network was trained on four inputs, presented in a random order, each simulating an object. Inputs were generated using a custom Python code (available here: https://github.com/frdarya/CompModel together with all necessary code to reproduce the data reported here). Out of 54 feature dimensions, every object had 12 active units (clamped to 1) and three units clamped to 0.9, representing a previously presented cue (Schapiro et al., 2017). Two cues were randomly set, one for ‘man-made’ objects and one for ‘natural’ objects, simulating the conditions used in Frank and Montaldi (2019). Therefore, two stimuli were associated with a ‘man-made’ category and the other two with a ‘natural’ category. During a training trial (100 processing cycles), the network was presented with a single object and the associated cue for two minus phases and a single plus phase. A full set of trials including all four objects completed an epoch. The network was trained for 10 epochs and tested following the last one.

At test, no changes were applied to connection weights, keeping the network at a retrieval-like neutral state. In every test trial, we presented the network an input and recorded the activation levels of each hidden layer. We employed cue units to simulate the ‘expected’ and ‘unexpected’ conditions used in our previous study (Frank & Montaldi, 2019). Expected trials had the same cue-object association the network was trained on, whereas in unexpected trials the cue was flipped (i.e. a man-made cue for a natural object, and vice versa). Critically, because the network has learned the contingency of both cues, flipping them at test does not create a novel input, but rather an unexpected pairing of learned inputs. In addition to testing each encoded object, we also created parametrically manipulated similar foils (expected and unexpected). These foils were created pseudo-randomly by varying the percentage of overlap between the target (original item) and the foil, ensuring flipped units between foils were also independent (e.g. units changed in F50 were not the same as those changed in F67). The very high similarity foil (F85) had an 85% overlap with the target, F67 had 67% overlap with the target, F50 had 50% overlap, whereas the lowest similarity foil (F33) had only 33% overlap with the encoded target. Therefore, for each object, 10 test trials were used (one target and four foils, each tested once as expected and once as unexpected), resulting in 40 test trials altogether (See Figure1b for illustration of the stimuli used).

### Lesions

To examine the contribution of pattern separation to differences in representational dissimilarity between expected and unexpected inputs, we simulated lesions in TSP. The following projection strengths were set to 0: EC_in_ → DG, EC_in_ → CA3, DG → CA3, CA3 → CA3, and CA3 → CA1. These lesions will reveal the independent function of MSP to the representation created in CA1 without any pattern separation from DG and CA3 (as DG and CA3 are ‘turned off’, they cannot be examined). As noted by Schapiro et al. (2017) MSP lesions do not reveal the independent contribution of TSP as MSP serves communications between EC and TSP, therefore they were not examined.

### Analyses

To assess representational similarity between targets and foils in both expectation conditions, we recorded unit activity in each hidden layer, per test trial, and calculated distances as 1-Pearson correlations. Therefore, each batch had one 40×40 representational dissimilarity matrix (RDM), capturing all trials across conditions and objects. To examine differences between expected and unexpected trials, we averaged across objects and computed two 5×5 RDMs – one correlating expected-expected trials (EE), another unexpected-unexpected (UU; symmetrical matrices, meaningless diagonals). It is important to note this analysis overcomes the inherent difference in number of slots changed between expected and unexpected inputs. For example, the correlation between unexpected target and unexpected F85 is only driven by their perceptual similarity, as both of them had the same number of slots changed. Therefore, by comparing correlations from EE RDMs to ones from UU RDMs, we could examine how the perceptual similarity between inputs was modulated by the expectation manipulation. For comparisons between theoretical RDMs and simulated data we used the RSAtoolbox (Nili et al., 2014). Kendall’s Tau A was computed to assess the second-order correlation between categorical models and data RDMs; Pearson’s r was used for comparison between models of input similarity (i.e. multiple computational models) and data RDMs (Nili et al., 2014). All tests were FDR-corrected and analyses were done per batch and then averaged across them, using each randomly reinitialised network batch as a random effect.

## Author contribution

D.F., M.M. and D.M. designed the paradigm. D.F. built the model and analysed the data. D.F., M.M. and D.M. discussed the results and wrote the manuscript.

## References

Aisa, B., Mingus, B., & O’Reilly, R. (2008). The Emergent neural modeling system. Neural Networks, 21(8), 1146–1152. https://doi.org/10.1016/j.neunet.2008.06.016

Axmacher, N., Cohen, M. X., Fell, J., Haupt, S., Dümpelmann, M., Elger, C. E., … Ranganath, C. (2010). Intracranial EEG Correlates of Expectancy and Memory Formation in the Human Hippocampus and Nucleus Accumbens. Neuron, 65(4), 541–549. https://doi.org/10.1016/j.neuron.2010.02.006

Axmacher, N., Mormann, F., Fernández, G., Elger, C. E., & Fell, J. (2006). Memory formation by neuronal synchronization. Brain Research Reviews, 52(1), 170–182. https://doi.org/10.1016/j.brainresrev.2006.01.007

Bar, M. (2009). The proactive brain : memory for predictions, 1235–1243. https://doi.org/10.1098/rstb.2008.0310

Chen, J., Olsen, R. K., Preston, A. R., Glover, G. H., & Wagner, A. D. (2011). Associative retrieval processes in the human medial temporal lobe: hippocampal retrieval success and CA1 mismatch detection. Learning & Memory, 18(8), 523–528. https://doi.org/10.1101/lm.2135211

Douchamps, V., Jeewajee, A., Blundell, P., Burgess, N., & Lever, C. (2013). Evidence for Encoding versus Retrieval Scheduling in the Hippocampus by Theta Phase and Acetylcholine. Journal of Neuroscience, 33(20), 8689–8704. https://doi.org/10.1523/JNEUROSCI.4483-12.2013

Elfman, K. W., Aly, M., & Yonelinas, A. P. (2014). Neurocomputational account of memory and perception: Thresholded and graded signals in the hippocampus. Hippocampus, 24(12), 1672–1686. https://doi.org/10.1002/hipo.22345

Fitch, T. E., Sahr, R. N., Eastwood, B. J., Zhou, F. C., & Yang, C. R. (2006). Dopamine D1/5 Receptor Modulation of Firing Rate and Bidirectional Theta Burst Firing in Medial Septal/Vertical Limb of Diagonal Band Neurons In Vivo. Journal of Neurophysiology, 95(5), 2808–2820. https://doi.org/10.1152/jn.01210.2005

Frank, D., Gray, O., & Montaldi, D. (2019). SOLID-Similar object and lure image database. Behavior Research Methods. https://doi.org/10.3758/s13428-019-01211-7

Frank, D., & Montaldi, D. (2019). Pattern separation is the key driver of expectation-modulated memory. bioRxiv, 577791. https://doi.org/10.1101/577791

Gluck, M. a, Meeter, M., & Myers, C. E. (2003). Computational models of the hippocampal region: linking incremental learning and episodic memory. Trends in Cognitive Sciences, 7(6), 269–276. https://doi.org/10.1016/S1364-6613(03)00105-0

Gruber, M. J., Hsieh, L.-T., Staresina, B. P., Elger, C. E., Fell, J., Axmacher, N., & Ranganath, C. (2018). Theta Phase Synchronization between the Human Hippocampus and Prefrontal Cortex Increases during Encoding of Unexpected Information: A Case Study. Journal of Cognitive Neuroscience, 30(11), 1646–1656. https://doi.org/10.1162/jocn_a_01302

Hanslmayr, S., Staresina, B. P., & Bowman, H. (2016). Oscillations and Episodic Memory: Addressing the Synchronization/Desynchronization Conundrum. Trends in Neurosciences, 39(1), 16–25. https://doi.org/10.1016/j.tins.2015.11.004

Hasselmo, M. E., Bodelon, C., & Wyble, B. P. (2002). A Proposed Function for Hippocampal Theta Rhythm?: Separate Phases of Encoding and Retrieval Enhance Reversal of Prior Learning. Neural Computation, 14, 793–817.

Hulbert, J. C., & Norman, K. A. (2015). Neural Differentiation Tracks Improved Recall of Competing Memories Following Interleaved Study and Retrieval Practice. Cerebral Cortex, 25(10), 3994–4008. https://doi.org/10.1093/cercor/bhu284

Kafkas, A., & Montaldi, D. (2015). Striatal and midbrain connectivity with the hippocampus selectively boosts memory for contextual novelty. Hippocampus, 12, 1–12. https://doi.org/10.1002/hipo.22434

Kafkas, A., & Montaldi, D. (2018a). Expectation affects learning and modulates memory experience at retrieval. Cognition, 180(July), 123–134. https://doi.org/10.1016/j.cognition.2018.07.010

Kafkas, A., & Montaldi, D. (2018b). How do memory systems detect and respond to novelty? Neuroscience Letters, (January), 0–1. https://doi.org/10.1016/j.neulet.2018.01.053

Ketz, N., Morkonda, S. G., & O’Reilly, R. C. (2013). Theta Coordinated Error-Driven Learning in the Hippocampus. PLoS Computational Biology, 9(6), e1003067. https://doi.org/10.1371/journal.pcbi.1003067

Kriegeskorte, N., Mur, M., & Bandettini, P. (2008). Representational similarity analysis – connecting the branches of systems neuroscience. Frontiers in Systems Neuroscience, 84(4), 368–374. https://doi.org/10.3389/neuro.06.004.2008

Kumaran, D., & Maguire, E. a. (2007). Match Mismatch Processes Underlie Human Hippocampal Responses to Associative Novelty. Journal of Neuroscience, 27(32), 8517–8524. https://doi.org/10.1523/JNEUROSCI.1677-07.2007

Kunec, S., Hasselmo, M. E., & Kopell, N. (2005). Encoding and Retrieval in the CA3 Region of the Hippocampus: A Model of Theta-Phase Separation. Journal of Neurophysiology, 94(1), 70–82. https://doi.org/10.1152/jn.00731.2004

Leutgeb, J. K., Leutgeb, S., Moser, M., & Moser, E. I. (2007). Pattern Separation in the Dentate Gyrus and CA3 of the Hippocampus, 315(February), 961–966.

Lisman, J. E., & Grace, A. a. (2005). The Hippocampal-VTA Loop: Controlling the Entry of Information into Long-Term Memory. Neuron, 46(5), 703–713. https://doi.org/10.1016/j.neuron.2005.05.002

Long, N. M., Lee, H., & Kuhl, B. a. (2016). Hippocampal Mismatch Signals Are Modulated by the Strength of Neural Predictions and Their Similarity to Outcomes. The Journal of Neuroscience, 36(50), 12677–12687. https://doi.org/10.1523/JNEUROSCI.1850-16.2016

Marr, D. (1971). Simple memory: a theory for archicortex. Philosophical Transactions of the Royal Society B: Biological Sciences, 262, 23–81.

McClelland, J. L., McNaughton, B. L., & O’Reilly, R. C. (1995). Why there are complementary learning systems in the hippocampus and neocortex: Psychological Review, 102(3), 419–457. https://doi.org/7624455

Meeter, M., Murre, J. M. J., & Talamini, L. M. (2004). Mode shifting between storage and recall based on novelty detection in oscillating hippocampal circuits. Hippocampus, 14(6), 722–741. https://doi.org/10.1002/hipo.10214

Mizumori, S. J. Y. (2016). Self Regulation of Memory Processing Centers of the Brain, 199–225. https://doi.org/10.1007/978-3-319-15759-7

Nili, H., Wingfield, C., Walther, A., Su, L., Marslen-Wilson, W., & Kriegeskorte, N. (2014). A Toolbox for Representational Similarity Analysis. PLoS Computational Biology, 10(4). https://doi.org/10.1371/journal.pcbi.1003553

Norman, K. A., & O’Reilly, R. C. (2003). Modeling hippocampal and neocortical contributions to recognition memory: A complementary-learning-systems approach. Psychological Review, 110(4), 611–646. https://doi.org/10.1037/0033-295X.110.4.611

O’Reilly, R. C., & Munakata, Y. (2000). Computational explorations in cognitive neuroscience: Understanding the mind by simulating the brain. Cambridge, MA: MIT Press.

Pilly, P. K., Howard, M. D., & Bhattacharyya, R. (2018). Modeling Contextual Modulation of Memory Associations in the Hippocampus. Frontiers in Human Neuroscience, 12(November), 442. https://doi.org/10.3389/fnhum.2018.00442

Pine, A., Sadeh, N., Ben-Yakov, A., Dudai, Y., & Mendelsohn, A. (2018). Knowledge acquisition is governed by striatal prediction errors. Nature Communications, 9(1), 1–14. https://doi.org/10.1038/s41467-018-03992-5

Schapiro, A. C., Turk-Browne, N. B., Botvinick, M. M., & Norman, K. A. (2017). Complementary learning systems within the hippocampus: a neural network modelling approach to reconciling episodic memory with statistical learning. Philosophical Transactions of the Royal Society B: Biological Sciences, 372(1711), 20160049. https://doi.org/10.1098/rstb.2016.0049

Shohamy, D., & Wagner, A. D. (2008). Integrating Memories in the Human Brain: Hippocampal-Midbrain Encoding of Overlapping Events. Neuron, 60(2), 378–389. https://doi.org/10.1016/j.neuron.2008.09.023

Vago, D. R., Bevan, A., & Kesner, R. P. (2007). The role of the direct perforant path input to the CA1 subregion of the dorsal hippocampus in memory retention and retrieval. Hippocampus, 17(10), 977–987. https://doi.org/10.1002/hipo.20329

Valenti, O., Mikus, N., & Klausberger, T. (2018). The cognitive nuances of surprising events: exposure to unexpected stimuli elicits firing variations in neurons of the dorsal CA1 hippocampus. Brain Structure and Function, 0(0), 1–29. https://doi.org/10.1007/s00429-018-1681-6

Xue, G. (2018). The Neural Representations Underlying Human Episodic Memory. Trends in Cognitive Sciences, 22(6), 544–561. https://doi.org/10.1016/j.tics.2018.03.004

Yassa, M. a, & Stark, C. E. L. (2011). Pattern separation in the hippocampus. Trends in Neurosciences, 34(10), 515–525. https://doi.org/10.1016/j.tins.2011.06.006

